# The two Gtsf paralogs in silkworms orthogonally activate their partner PIWI proteins for target cleavage

**DOI:** 10.1101/2022.06.30.498350

**Authors:** Natsuko Izumi, Keisuke Shoji, Takashi Kiuchi, Susumu Katsuma, Yukihide Tomari

**Affiliations:** Laboratory of RNA Function, Institute for Quantitative Biosciences, The University of Tokyo, Bunkyo-ku, Tokyo 113-0032, Japan; Department of Agricultural and Environmental Biology, Graduate School of Agricultural and Life Sciences, The University of Tokyo, Bunkyo-ku, Tokyo 113-8657, Japan

**Keywords:** *Bombyx mori*, Gtsf, piRNA, PIWI

## Abstract

The PIWI-interacting RNA (piRNA) pathway is a protection mechanism against transposons in animal germ cells. Most PIWI proteins possess piRNA-guided endonuclease activity, which is critical for silencing transposons and producing new piRNAs. Gametocyte-specific factor 1 (Gtsf1), an evolutionarily conserved zinc finger protein, promotes catalysis by PIWI proteins. Many animals have multiple Gtsf1 paralogs; however, their respective roles in the piRNA pathway are not fully understood. Here, we dissected the roles of Gtsf1 and its paralog Gtsf1-like (Gtsf1L) in the silkworm piRNA pathway. We found that Gtsf1 and Gtsf1L preferentially bind the two silkworm PIWI paralogs, Siwi and BmAgo3, respectively, and facilitate the endonuclease activity of each PIWI protein. This orthogonal activation effect was further supported by specific reduction of BmAgo3-bound *Masculinizer* piRNA and Siwi-bound *Feminizer* piRNA, the unique piRNA pair required for silkworm feminization, upon depletion of Gtsf1 and Gtsf1L, respectively. Our results indicate that the two Gtsf paralogs in silkworms activate their respective PIWI partners, thereby facilitating the amplification of piRNAs.

## Introduction

PIWI-interacting RNAs (piRNAs) play a central role in transposon silencing in animal germ cells, thereby maintaining their genome integrity and ensuring their proper development. piRNAs are ∼24−31 nt in length and guide PIWI-clade Argonaute (Ago) proteins to complementary targets, repressing their expression at the transcriptional and post-transcriptional levels (Ozata *et al*, 2019; Iwasaki *et al*, 2015). Most PIWIs possess the endonuclease activity called “slicer” and can directly cleave target RNAs (Reuter *et al*, 2011; De Fazio *et al*, 2011; Brennecke *et al*, 2007; Gunawardane *et al*, 2007). PIWI-catalyzed RNA cleavage not only silences target transposons but also produces new piRNAs (Ozata *et al*, 2019); target cleavage by a piRNA-guided PIWI protein (e.g., Siwi in silkworms; Aubergine [Aub] in flies) produces a long precursor of a new complementary piRNA (called pre-pre-piRNA), which is loaded into another PIWI protein (e.g., BmAgo3 in silkworms; Ago3 in flies) with its 5’ end anchored. PIWI-loaded pre-pre-piRNA is then cleaved at a downstream position by the endonuclease Zucchini (Zuc) or by another piRNA-guided PIWI protein, resulting in the production of a shorter precursor (called pre-piRNA) (Izumi *et al*, 2020; Gainetdinov *et al*, 2018; Ozata *et al*, 2019). The 3’ end of pre-piRNA is further shortened by the exonuclease Trimmer (PNLDC1 in mice) and methylated by the 2’-*O*-methyltransferase Hen1 (HENMT1 in mice) to generate a mature piRNA (Nishimura *et al*, 2018; Ding *et al*, 2017; Izumi *et al*, 2016; Zhang *et al*, 2017; Horwich *et al*, 2007; Saito *et al*, 2007; Kirino & Mourelatos, 2007). This piRNA biogenesis pathway, which depends on reciprocal target cleavage by a pair of PIWI proteins, is called the ping-pong cycle. Since PIWI proteins cleave target RNAs at the specific position across the 10th and 11th nucleotides of the guiding piRNA, piRNA pairs produced by the ping-pong cycle show a 10-nt complementary overlap at their 5’ ends. In general, “ping” PIWI proteins (e.g., Siwi in silkworms; Aub in flies) preferentially bind piRNAs with 5’ uracil (1U), and their partner “pong” PIWI proteins (e.g., BmAgo3 in silkworms; Ago3 in flies) are loaded with piRNAs bearing adenine at the 10th nucleotide (10A) through the ping-pong cycle (Brennecke *et al*, 2007; Gunawardane *et al*, 2007; Kawaoka *et al*, 2009; Nishida *et al*, 2015; Wang *et al*, 2014). Coupled with the ping-pong cycle, the 3’ fragment of the cleavage products by Zuc is loaded into the next PIWI protein (e.g., Siwi in silkworms; Piwi and Aub in flies) as a new pre-pre-piRNA, which is then cleaved at a downstream position again and processed into a mature piRNA. As a result, a series of “trailing” piRNAs are consecutively produced in the downstream region (Gainetdinov *et al*, 2018; Ozata *et al*, 2019; Mohn *et al*, 2015; Han *et al*, 2015). PIWI proteins and other piRNA biogenesis factors often localize to perinuclear membrane-less granules (called “nuage”), processing bodies (P-bodies), and/or mitochondrial surfaces (Ozata *et al*, 2019; Iwasaki *et al*, 2015). Some factors shuttle between different compartments during piRNA biogenesis (Ge *et al*, 2019), and dynamic subcellular partitioning of piRNA factors is important for the fidelity of piRNA production (Chung *et al*, 2021).

Gametocyte-specific factor 1 (Gtsf1), an evolutionarily conserved small zinc finger protein, has been characterized as an essential piRNA factor in mice and flies (Ipsaro & Joshua-Tor, 2022). The *Drosophila* homolog of Gtsf1, Asterix, binds Piwi and functions in Piwi-mediated transcriptional silencing in the nucleus, while *asterix* mutant has no apparent effects on the piRNA biogenesis (Muerdter *et al*, 2013; Ohtani *et al*, 2013; Dönertas *et al*, 2013). On the other hand, mouse GTSF1 is required for the production of MIWI2-bound piRNAs, and *Gtsf1* knockout (KO) exhibits similar phenotypes as *Miwi2* KO (Yoshimura *et al*, 2009, 2018). In silkworms, loss of Gtsf1 causes defects in gametogenesis, sex determination, and transposon silencing accompanied by downregulation of piRNAs (Chen *et al*, 2020). Recently, it was reported that mouse GTSF1 potentiates the otherwise poor endonucleolytic activity of MIWI and MILI, and that silkworm Gtsf1 has a similar activity for Siwi but not for BmAgo3 (Arif *et al*, 2022). Considering that sex determination in silkworms is mediated by a unique Siwi-bound piRNA called *Feminizer* (*Fem*) piRNA (Kiuchi *et al*, 2014), the phenotypes observed in silkworm *Gtsf1* mutant may be explained by the dysfunction of Siwi catalysis.

Many animals possess multiple Gtsf paralogs; flies have four (Gtsf1/Asterix, CG14036, CG32625, and CG34283), mice have three (GTSF1, GTSF1L, and GTSF2), and silkworms have two (Gtsf1 and Gtsf1-like [Gtsf1L]) (Fig EV1A). However, their respective roles in the piRNA pathway are not fully understood. Here, we dissected the role of the two Gtsf paralogs in silkworms in the piRNA pathway, using a silkworm ovarian cell line called BmN4 and silkworm embryos. We found that Gtsf1 and Gtsf1L preferentially bind the two silkworm PIWI proteins, Siwi and BmAgo3, respectively, and promote the slicer activity of each PIWI protein. Their orthogonal action on the partner PIWI protein was further supported by specific reduction of *Masculinizer* (*Masc*) and *Fem* piRNAs, the ping-pong pair required for silkworm sex determination, in depletion of Gtsf1 and Gtsf1L, respectively. These results suggest that the two Gtsf paralogs in silkworms independently function in the ping-pong cycle by facilitating the slicer activity of the respective partner PIWI proteins.

## Results and Discussion

### Gtsf1 and Gtsf1L bind and activate Siwi and BmAgo3, respectively

Silkworms have two Gtsf1 paralogs, Gtsf1 and Gtsf1-like (Gtsf1L), which have the characteristic tandem CHHC-type zinc-finger domains and show 27.8% similarity (Fig EV1B). We have recently reported that silkworm Gtsf1 promotes the slicer activity of Siwi but not that of BmAgo3 (Arif *et al*, 2022). Consistent with this, physical interaction between Gtsf1 and Siwi has been demonstrated (Chen *et al*, 2020). On the other hand, the function of Gtsf1L remains unclear. To see if Gtsf1L binds any PIWI protein(s), we transiently expressed epitope-tagged Gtsf1 or Gtsf1L in BmN4 cells and examined the interaction with endogenous Siwi and BmAgo3. Gtsf1 was preferentially immunoprecipitated with Siwi, while Gtsf1L specifically interacted with BmAgo3 (Fig 1A). This orthogonal interaction was also supported by AlphaFold-Multimer (Fig EV1C and D), and the C-terminal structure of Gtsf1 and Gtsf1L was predicted with higher reliability when complexed with Siwi and BmAgo3, respectively (Fig EV1E and F). As with mouse GTSF1, which directly interacts with TDRD9 [Spindle-E (SpnE) ortholog in mice] (Yoshimura *et al*, 2018), SpnE was also co-immunoprecipitated with Gtsf1 but not with Gtsf1L (Figs 1A and EV1G).

**Figure 1.**
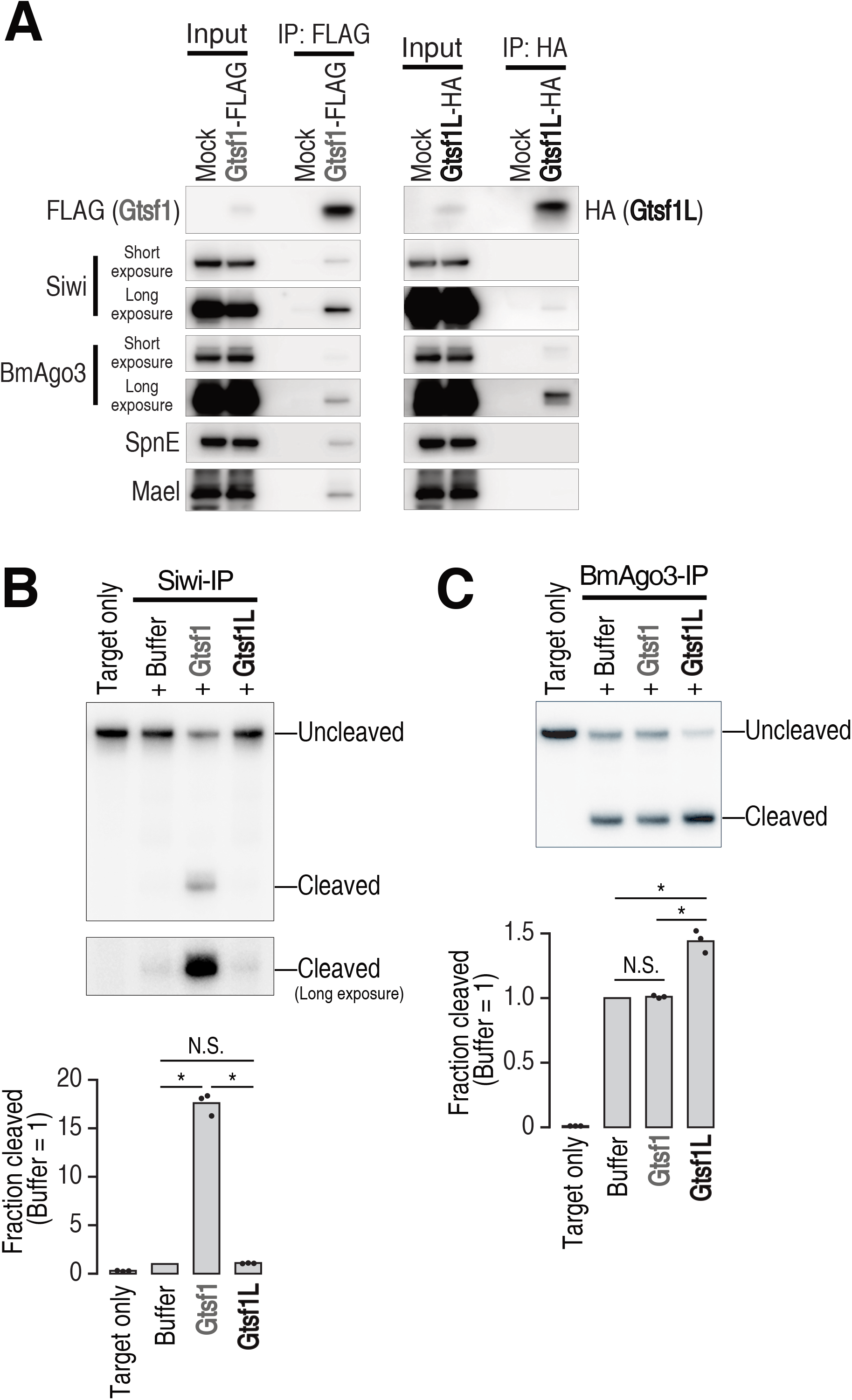
Gtsf1 and Gtsf1L bind Siwi and BmAgo3, respectively, and activate each PIWI protein. **A** Western blot analysis of immunoprecipitated tagged-Gtsf1 and -Gtsf1L from BmN4 cells. **B, C** *In vitro* target cleavage assay of immunoprecipitated Siwi (B) or BmAgo3 (C) from BmN4 cells. Target RNA against an abundant endogenous Siwi- (B) or BmAgo3-bound (C) piRNA was used. The graphs show the fraction cleaved normalized to that of the buffer-only sample. The values are means of three independent experiments. Dots, individual values. Paired t-test with Bonferroni adjusted *p*-values are 0.0046 (Buffer vs Gtsf1 in [B]), 0.0049 (Gtsf1 vs Gtsf1L in [B]), 0.04 (Buffer vs Gtsf1L in [C]), and 0.039 (Gtsf1 vs Gtsf1L in [C]). N.S., not significant.

Considering this different binding preference, we re-examined the effect of recombinant Gtsf proteins on the slicer activity of Siwi or BmAgo3 *in vitro*. We first prepared a target RNA complementary to an endogenous Siwi-bound piRNA and monitored target cleavage by Siwi immunoprecipitate in the presence or absence of recombinant Gtsf1 or Gtsf1L (Figs 1B and EV1H). As previously reported (Arif *et al*, 2022), we confirmed that Gtsf1 but not Gtsf1L promotes target cleavage by Siwi (Fig 1B). In the previous study, the target cleavage activity by BmAgo3 was extremely inefficient, making it difficult to conclusively examine the activity of Gtsf1L (Arif *et al*, 2022). We therefore searched for a better target sequence for an endogenous BmAgo3-bound piRNA. By using this new target, we could detect modest but significant enhancement of BmAgo3-mediated target cleavage by Gtsf1L but not by Gtsf1 (Fig 1C). Taken together, these data suggest that Gtsf1 and Gtsf1L specifically interact with Siwi and BmAgo3, respectively, and facilitate the slicer activity of each PIWI protein.

### Gtsf1 and Gtsf1L accumulate in the complex of catalytically inactive Siwi and BmAgo3, respectively

Because both Gtsf1 and Gtsf1L are expected to function in the catalytic process of PIWI proteins (Fig 1B and C) (Arif *et al*, 2022), we next examined their interaction with PIWI catalytic mutants. We co-expressed epitope-tagged Gtsf1 and wild-type (WT) or catalytically inactive (D670A) Siwi in BmN4 cells, and immunoprecipitated Gtsf1. Compared to Siwi-WT, Siwi-D670A showed markedly increased association with Gtsf1 (Fig 2A). Although Gtsf1 was broadly distributed in the nucleus and cytoplasm with little co-localization with Siwi-WT, Gtsf1 showed partial but clear co-localization with Siwi-D670A granules (Fig 2B), suggesting that Siwi-D670A traps Gtsf1. We previously reported that Siwi-D670A localizes to P-bodies with SpnE (Chung *et al*, 2021). In addition, in mouse gonocytes, GTSF1 localizes to piP-bodies (Yoshimura *et al*, 2018), which share components with P-bodies, together with MIWI2, TDRD9 (SpnE ortholog in mice) and MAEL (Aravin *et al*, 2009). Since Gtsf1 interacts with SpnE and Mael (Fig 1A), we expected that Gtsf1 trapped by Siwi-D670A is localized to P-bodies. Indeed, we observed granular co-localization among Siwi-D670A, Gtsf1, and Dcp2, a P-body marker (Fig 2C). In contrast, co-localization between Gtsf1 and Dcp2 was not detected when Siwi-WT was expressed (Fig 2C). Thus, Gtsf1 is not a stable resident of P-bodies but can be trapped in P-bodies by catalytically inactive Siwi.

**Figure 2.**
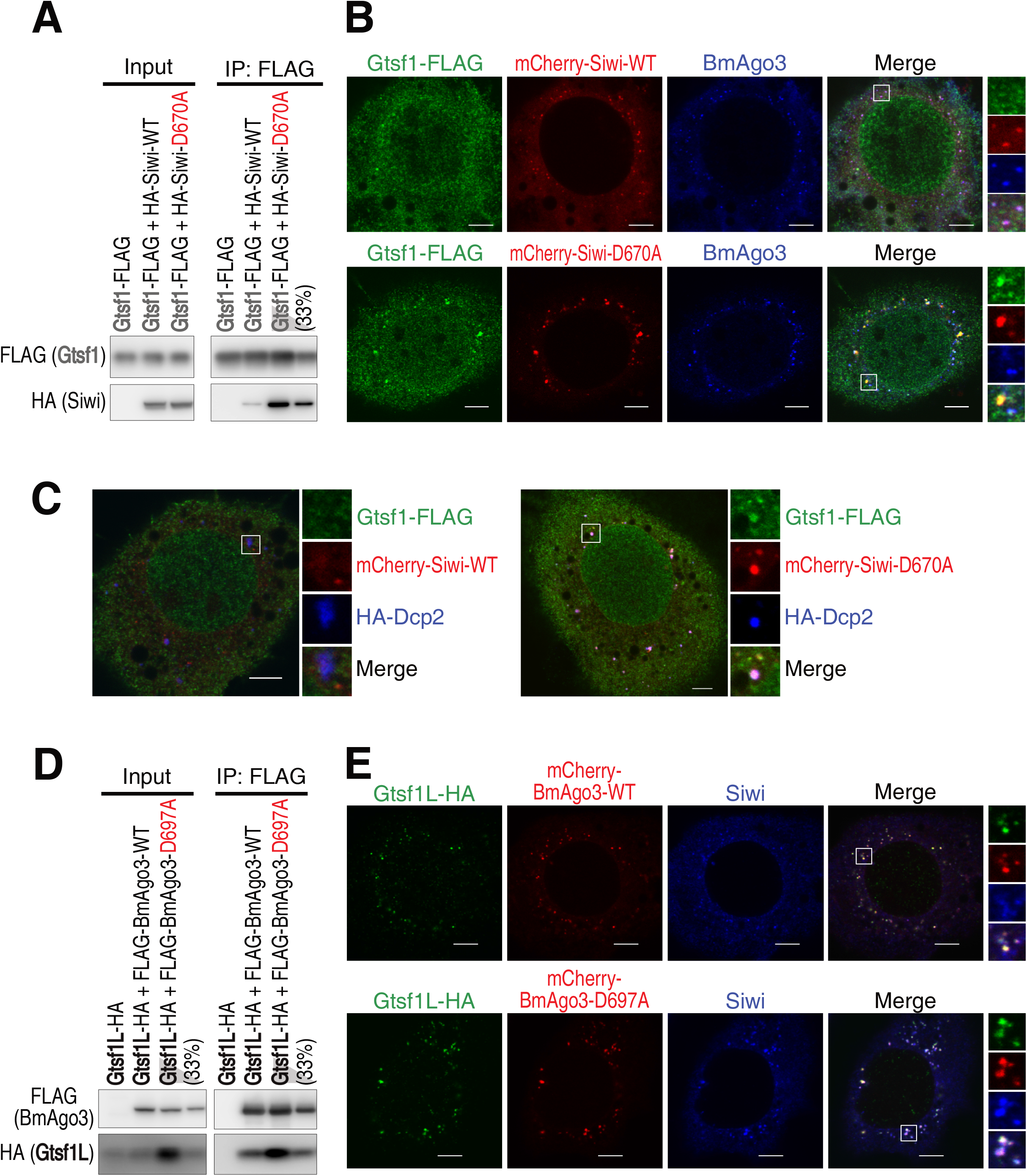
Gtsf1 and Gtsf1L show increased association with catalytically inactive Siwi and BmAgo3, respectively. **A** Western blot analysis of the interaction between Gtsf1-FLAG and HA-tagged wild-type (WT) Siwi or catalytically inactive Siwi-D670A. **B** Subcellular localization of Gtsf1-FLAG co-expressed with mCherry-Siwi or -Siwi-D670A. Scale bar, 5 μm. **C** Subcellular localization of Gtsf1-FLAG co-expressed with mCherry-Siwi or -Siwi-D670A, and HA-Dcp2, a P-body marker. Scale bar, 5 μm. **D** Western blot analysis of the interaction between Gtsf1L-HA and FLAG-tagged wild-type (WT) BmAgo3 or catalytically inactive BmAgo3-D697A. **E** Subcellular localization of Gtsf1L-HA co-expressed with mCherry-BmAgo3 or -BmAgo3-D697A. Scale bar, 5 μm.

We next investigated the association between Gtsf1L and the catalytic mutant of BmAgo3 (D697A). We co-expressed epitope-tagged BmAgo3-WT or -D697A and Gtsf1L in BmN4 cells, and immunoprecipitated BmAgo3. The steady-state level of Gtsf1L was greatly increased by the co-expression of BmAgo3-D697A (Fig 2D, input). Correspondingly, co-immunoprecipitation of Gtsf1L with BmAgo3-D697A was markedly higher than that with BmAgo3-WT (Fig 2D). Unlike Gtsf1, Gtsf1L was well co-localized with BmAgo3 granules regardless of the catalytic activity of BmAgo3 (Fig 2E). The catalytic mutation of BmAgo3 slightly intensified the BmAgo3-Gtsf1L granules, and also increased the granule formation of Siwi, a part of which partially overlapped with BmAgo3 and Gtsf1L (Fig 2E, BmAgo3-D697A).

We next examined the behavior of Gtsf1L in *Siwi* knockdown (KD), where the handover of RNAs from BmAgo3 to Siwi in the ping-pong cycle is inhibited (Nishida *et al*, 2020), thereby creating an analogous situation as the catalytic inactivation of BmAgo3. Similar to the overexpression of BmAgo3-D697A, *Siwi*-KD increased the steady-state abundance of Gtsf1L and its association with BmAgo3 (Fig EV2A). In agreement with this, Gtsf1L strongly accumulated in the enlarged BmAgo3 granules in *Siwi*-KD (Fig EV2B). In contrast to Gtsf1L, neither the Gtsf1-Siwi interaction nor their co-localization was changed by *BmAgo3*-KD (Fig EV2C and D). The different behavior of Gtsf1 and Gtsf1L in *BmAgo3*-KD or *Siwi*-KD may reflect the fact that Siwi, but not BmAgo3, can operate homotypic ping-pong, especially in the absence of the counterpart PIWI protein (N.I., K.S., T.K., S.K., and Y.T., unpublished observation), similarly to Aub:Aub homotypic ping-pong in *ago3* mutant flies (Li *et al*, 2009a). In sum, our observations that Gtsf1 and Gtsf1L accumulate in the complex of their catalytically inactive PIWI partner underscore their roles in the PIWI catalytic process.

### Loss of Gtsf1 and Gtsf1L reduces *Masc* and *Fem* piRNAs, respectively

In silkworms, the ping-pong cycle normally operates between Siwi and BmAgo3 (Kawaoka *et al*, 2009; Nishida *et al*, 2015). If Gtsf1 and Gtsf1L promote the slicer activity of Siwi and BmAgo3, respectively, loss of Gtsf1 and Gtsf1L should differently affect Siwi- and BmAgo3-bound piRNAs. We therefore sequenced small RNAs from BmN4 cells treated with dsRNA for *Gtsf1* or *Gtsf1L*. Either knockdown modestly decreased total mature piRNAs and the strength of the ping-pong signature, suggesting the impairment of the ping-pong cycle (Fig EV3A and B). Because Siwi- and BmAgo3-bound piRNAs have the 1U and 10A bias, respectively (Kawaoka *et al*, 2009; Nishida *et al*, 2015), we next examined the change in 1U (but not 10A) and 10A (but not 1U) piRNAs. *Gtsf1*-KD modestly decreased both 1U- and 10A-piRNAs, while *Gtsf1L*-KD caused more severe reduction of 1U-piRNAs than 10A-piRNAs (Fig 3A and B). The ping-pong piRNAs are generally mapped densely in discrete genomic loci and their production can be heavily influenced by neighboring piRNAs; the 3’ end processing of a piRNA is often mediated by another piRNA-guided cleavage in the downstream region (Hayashi *et al*, 2016; Izumi *et al*, 2020). Therefore, we focused on an isolated, well-defined ping-pong pair, *Fem* and *Masc* piRNAs, which determine femaleness in silkworms (Fig 3C). *Fem* piRNA, deriving from the W chromosome, is exclusively loaded into Siwi and cleaves *Masc* mRNA, leading to the reduction of Masc protein and the suppression of masculinization (Kiuchi *et al*, 2014). *Fem* piRNA-mediated cleavage of *Masc* mRNA produces BmAgo3-bound *Masc* piRNA, which in turn cleaves *Fem* RNA and amplifies *Fem* piRNA via the ping-pong cycle (Kiuchi *et al*, 2014). No other evident ping-pong pair is found on *Fem* RNA and *Masc* mRNA, thus the effect from neighboring piRNAs can be ignored. Strikingly, *Gtsf1*-KD and *Gtsf1L*-KD specifically decreased *Masc* piRNA and *Fem* piRNA, respectively (Fig 3C). This data supports the idea that Gtsf1 activates Siwi to promote the production of BmAgo3-bound *Masc* piRNA, while Gtsf1L activates BmAgo3 to promote the production of Siwi-bound *Fem* piRNA.

**Figure 3.**
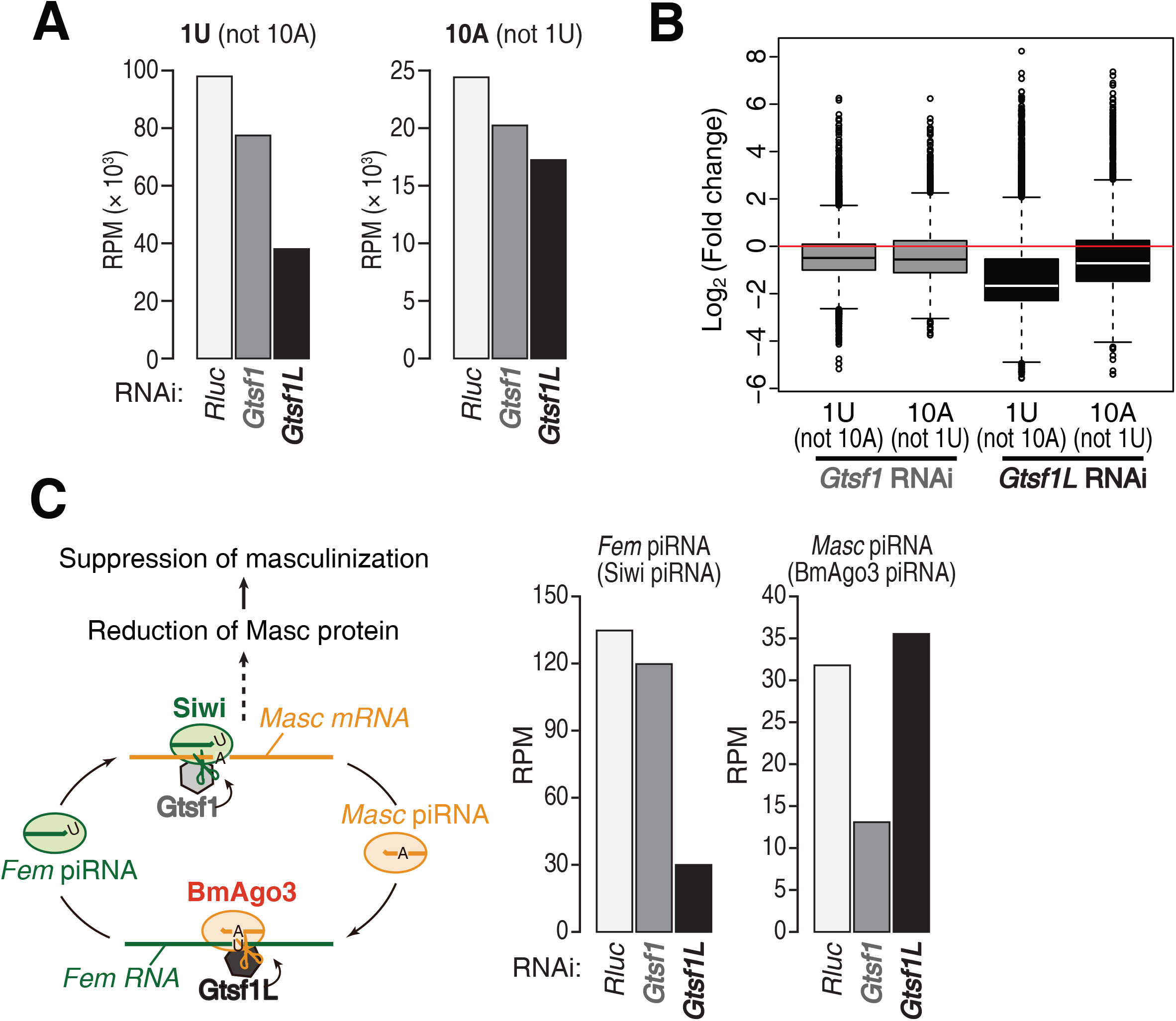
Knockdown of *Gtsf1* and *Gtsf1L* decreases *Masc* and *Fem* piRNAs, respectively. **A** Normalized reads of 1U (but not 10A) or 10A (but not 1U) small RNA from BmN4 cells transfected with dsRNA for *Renilla luciferase* (*Rluc*, control), *Gtsf1* or *Gtsf1L*. **B** Box plots showing the expression change of 1U (but not 10A) and 10A (but not 1U) piRNAs by knockdown of *Gtsf1* or *Gtsf1L* relative to control knockdown in BmN4 cells. Center line, median; box limits, upper and lower quartiles; whiskers, 1.5 × interquartile range; points, outliers. **C** Schematic representation of the production of *Fem* and *Masc* piRNAs via the ping-pong cycle (left). The expression of *Fem* and *Masc* piRNAs in *Gtsf1*- or *Gtsf1L*-knockdown (right).

### Gtsf1 and Gtsf1L independently function in the ping-pong cycle *in vivo*

In the early embryonic stages of silkworms, maternally inherited piRNAs initiate acute and massive amplification of new piRNAs via the ping-pong cycle (Kawaoka *et al*, 2011). In this regard, silkworm embryos may be better suited to monitor the effect of Gtsf proteins in the ping-pong cycle, compared to the cell line BmN4, where piRNAs are constantly produced and influencing each other in a steady state. To see if Gtsf1 and Gtsf1L function in a PIWI-type specific manner *in vivo*, we injected siRNAs against *Gtsf1* or *Gtsf1L* into silkworm embryos (Fig EV4A) and examined piRNA production after 120 hours post-injection. We prepared small RNA libraries from the embryos with two biological replicates per each knockdown condition. Consistent with the result in BmN4 cells, *Gtsf1*-KD and *Gtsf1L*-KD reduced the strength of the ping-pong signature (Fig EV4B). Moreover, *Masc* and *Fem* piRNA were specifically decreased by *Gtsf1*-KD and *Gtsf1L*-KD, respectively (Fig 4A), confirming the orthogonal functions of the two Gtsf paralogs *in vivo*. We also observed that, in general, 1U (but not 10A) piRNAs are downregulated by *Gtsf1L*-KD, whereas 10A (but not 1U) piRNAs are decreased by *Gtsf1*-KD (Fig 4B and C). To analyze this effect more precisely, we focused on 775 transposons in which 1U (but not 10A) piRNAs are disproportionally enriched on either of the two strands (“1U strand”) while 10A (but not 1U) piRNAs are reciprocally enriched on the other strand (“10A strand”). We then calculated the ratio of mapped piRNA reads on the 1U and 10A strands (“1U/10A strand bias”) for each transposon and monitored the change of the bias (Z-score) upon *Gtsf1*-KD or *Gtsf1L*-KD (Fig 4D). We found that, for nearly all the transposons analyzed, *Gtsf1*-KD increased the 1U/10A strand bias (i.e., 10A [but not 1U] piRNAs were more reduced than 1U [but not 10A] piRNAs), while *Gtsf1L*-KD decreased the 1U/10A strand bias (i.e., 1U [but not 10A] piRNAs were more downregulated). These results strengthen the idea that Gtsf1 and Gtsf1L orthogonally activate the target cleavage by Siwi and BmAgo3, thereby promoting the production of BmAgo3-bound and Siwi-bound piRNAs, respectively, *in vivo*.

**Figure 4.**
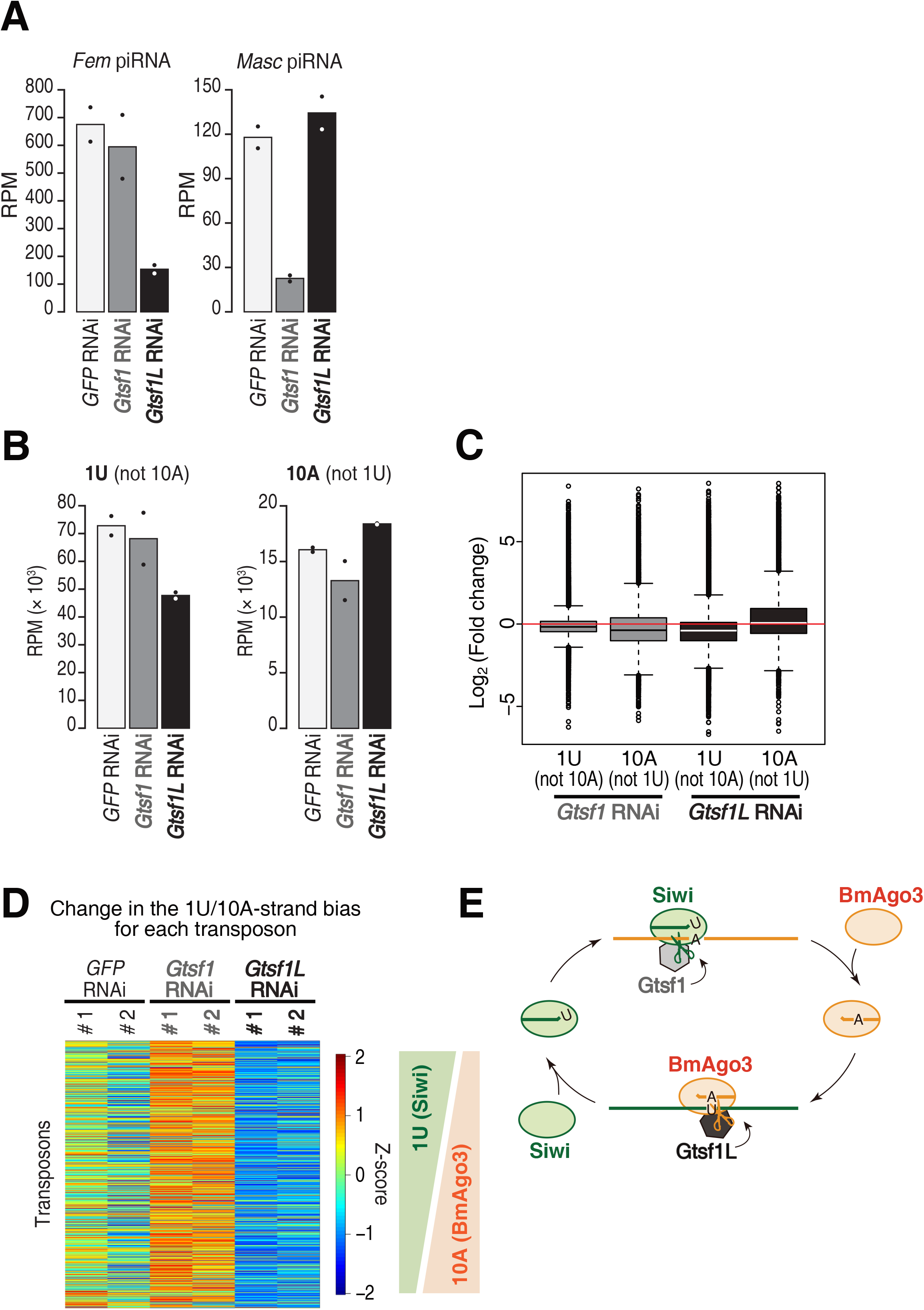
Gtsf1 and Gtsf1L separately function in the ping-pong cycle *in vivo*. **A** The expression of *Fem* and *Masc* piRNAs in silkworm embryos injected with siRNAs for *GFP* (control), *Gtsf1* or *Gtsf1L*. The values are means of two biological replicates. Dots, individual values. **B** Normalized reads of 1U (but not 10A) or 10A (but not 1U) small RNA from silkworm embryos injected with siRNAs for *GFP* (control), *Gtsf1* or *Gtsf1L*. The values are means of two biological replicates. Dots, individual values. **C** Box plots showing the expression change of 1U (but not 10A) and 10A (but not 1U) piRNAs by knockdown of *Gtsf1*or *Gtsf1L* relative to *GFP* (control) knockdown in embryos. Center line, median; box limits, upper and lower quartiles; whiskers, 1.5 × interquartile range; points, outliers. **D** Heatmap representation of relative changes in the 1U/10A strand bias between 6 samples for each transposon. The 1U/10A strand bias tends to increase (i.e., relative reduction of 10A-biased BmAgo3 piRNA) and decrease (i.e., relative reduction of 1U-biased Siwi piRNA) in *Gtsf1*- and *Gtsf1L*-knockdown, respectively. Two biological replicates per each knockdown condition. **E** A model for the role of two silkworm Gtsf proteins in the ping-pong cycle. Gtsf1 and Gtsf1L contribute to the production of BmAgo3- and Siwi-piRNAs by facilitating target cleavage of Siwi and BmAgo3, respectively.

In this study, we revealed that the two Gtsf paralogs in silkworms, Gtsf1 and Gtsf1L, individually function in the ping-pong cycle by promoting the slicer activity of Siwi and BmAgo3, respectively (Fig 4E). In mice, GTSF1 accelerates the target cleavage by MILI and MIWI, while the other two GTSF paralogs, GTSF1L and GTSF2, can potentiate the catalytic activity of MIWI, at least *in vitro* (Arif *et al*, 2022). However, double knockout mice of *Gtsf1l* and *Gtsf2* exhibit no defects in spermatogenesis and retrotransposon silencing (Takemoto *et al*, 2016), suggesting that GTSF1 alone is shared by MIWI and MILI for their catalytic activation in mice. On the other hand, we found that the two Gtsf proteins in silkworms have specific preferences for the two PIWI proteins, respectively, and cannot compensate for each other. Thus, silkworm Gtsf proteins have evolved to specialize for each PIWI protein. Both Gtsf1 and Gtsf1L orthologs are found in lepidoptera and mosquitos, therefore their separate roles in the ping-pong cycle may be widely conserved in these insects. Sequence comparison of Gtsf homologs in multiple species has suggested that the diversity in their C-terminal unstructured region is involved in the PIWI specificity (Arif *et al*, 2022). In agreement with this hypothesis, the C-terminal structures of Gtsf1 and Gtsf1L were predicted with higher confidence when complexed with Siwi and BmAgo3, respectively, by AlphaFold-Multimer (Fig EV1E). Considering that the steady state abundance of Gtsf1L is markedly increased by co-expression of catalytically inactive BmAgo3 (Fig 2D), binding to BmAgo3 may stabilize the Gtsf1L protein itself.

How do Gtsf proteins accelerate target cleavage by PIWI? A recent structural study in *Ephydatia fluviatilis* PIWI revealed that after recognition of target RNA, extensive base pairing between piRNA and target RNA induces a conformational change in PIWI (Anzelon *et al*, 2021). Accordingly, it has been proposed that such a catalytic competent state of PIWI is stabilized by Gtsf1 (Arif *et al*, 2022). Indeed, AlphaFold-Multimer predicts that both Gtsf proteins locate around the catalytic PIWI domain (Fig EV1F). Moreover, the first zinc finger motif of mouse GTSF1, which has RNA binding activity, is required to promote PIWI slicing (Ipsaro *et al*, 2021; Arif *et al*, 2022). Therefore, it is likely that Gtsf binds to the PIWI domain and the target RNA simultaneously to stabilize the catalytic competent conformation of PIWI proteins. The observation that both silkworm Gtsf1 and Gtsf1L accumulate in the catalytically inactive PIWI complex also supports the idea that Gtsf proteins have a higher affinity to the target-bound state of PIWI proteins. Future structural analysis of the Gtsf-PIWI complex is awaited to understand how Gtsf binding contributes to the structural change of PIWIs during the target cleavage.

## Data availability

The sequencing data is deposited at the DDBJ database (https://www.ddbj.nig.ac.jp) under accession number DRA014298.

## Acknowledgements

Illumina sequencing was performed in the Vincent J. Coates Genomics Sequencing Laboratory at UC Berkeley, supported by NIH S10 OD018174 Instrumentation Grant. We thank P. Zamore and all the members of the Tomari laboratory for critical comments on the manuscript. This work was in part supported by a Grant-in-Aid for Scientific Research (S) (grant 18H05271 to Y.T.), and Grant-in-Aid for Scientific Research on Innovative Areas (grant 17H06431 to S.K. and T.K.), Grant-in-Aid for Scientific Research (C) (grant 19K06484 to N.I.) and Grant-in-Aid for Early-Career Scientists (grant 22K15082 to K.S.).

## Author Contributions

N.I., K.S. and Y.T. conceived and designed the experiments and wrote the manuscript. N.I. performed biochemical experiments and immunofluorescent staining. K.S. performed bioinformatics analyses. T.K. and S.K. performed siRNA injection experiments on silkworm embryos. Y.T. supervised the research. All the authors discussed the results and approved the manuscript.

## Conflict of interest

The authors declare that they have no conflict of interest.

## Materials and Methods

### Cell culture, plasmid transfection, and dsRNA transfection in BmN4 cells

BmN4 cells were cultured at 27 °C in IPL-41 medium (AppliChem) supplemented with 10% fetal bovine serum. For immunoprecipitation experiments, a total of 5–7.5 μg of plasmid and dsRNAs were transfected into BmN4 cells (2–2.5 × 10^6^ cells per 10 cm dish) with X-tremeGENE HP DNA Transfection Reagent (Sigma) and the transfected cells were harvested after 4 days. For co-transfection experiments, the second transfection was performed 2 days after the first transfection and the cells were harvested after an additional 4 days.. For immunofluorescence experiments, a total of 0.4 μg of plasmid and dsRNAs were transfected into BmN4 cells (4–6 × 10^4^ cells per glass bottom 35 mm dish) with X-tremeGENE HP DNA Transfection Reagent (Sigma) and cells were fixed 4 days later. For Fig EV2, transfection was repeated 2 days later and cells were fixed 4 days after the second transfection. For library preparation, 5 μg of dsRNAs were transfected into BmN4 cells (8 × 10^5^ cells per 10 cm dish) with X-tremeGENE HP DNA Transfection Reagent (Sigma) every 3 days four times. dsRNA preparation was described previously (Izumi *et al*, 2020). Template DNAs were prepared by PCR using primers containing T7 promoter listed in Table EV1.

### Plasmid construction

pIZ-FLAG-BmAgo3, pIZ-mCherry-Siwi, pIZ-mCherry-Siwi-D670A, pIZ-mCherry-BmAgo3, and pIZ-mCherry-BmAgo3-D697A were described previously (Kawaoka *et al*, 2009; Chung *et al*, 2021). The primer sequences for plasmid construction are listed in Table EV1.

#### pIExZ-Gtsf1-FLAG

cDNA fragment of Gtsf1 was amplified by RT-PCR from BmN4 total RNAs and cloned into pIExZ vector (Izumi *et al*, 2020) by In-fusion cloning kit (Takara).

#### pIZ-Gtsf1L-HA

A cDNA fragment of Gtsf1L was amplified by RT-PCR from BmN4 total RNAs and cloned into pIZ vector by In-fusion cloning kit (Takara).

#### pIZ-HA-Siwi, Siwi-D670A, pIZ-FLAG-BmAgo3-D697A

A DNA fragment coding HA-tagged Siwi was amplified by PCR and cloned into pIZ vector (Thermo Fisher/Invitrogen) by In-fusion cloning kit (Takara). pIZ-Siwi-D670A and pIZ-BmAgo3-D697A mutants were generated by site-directed mutagenesis.

#### pIExZ-HA-Dcp2

A DNA fragment coding HA-tagged Dcp2 was amplified by PCR and cloned into pIExZ vector (Izumi *et al*, 2020) by In-fusion cloning kit (Takara).

#### pCold-Gtsf1-7His, pCold-Gtsf1L-6His

A DNA fragment coding C-terminally His-tagged Gtsf1 or Gtsf1L was amplified by PCR and cloned into pCold vector (Takara) by In-fusion cloning kit (Takara). For Gtsf1L, the coding sequence was codon-optimized to *Bombyx mori* using EMBOSS Backtranseq (http://www.ebi.ac.uk/Tools/st/emboss_backtranseq/), and a synthesized DNA fragment (GenScript) was used as PCR template.

### Antibodies and Western blotting

Rabbit anti-Siwi, anti-BmAgo3, and anti-Mael antibodies were described previously (Chung *et al*, 2021; Izumi *et al*, 2020). Rabbit anti-SpnE antibody was generated by immunizing N-terminally His tagged recombinant SpnE (aa 2–124). Anti-FLAG (M2, Sigma), anti-HA (3F10, Roche), anti-α-Tubulin (B-5-1-2, Sigma) antibodies were purchased. For Fig EV2, the anti-BmAgo3 antibody was biotinylated using Biotin Labeling Kit-NH_2_ (DOJINDO). Chemiluminescence was induced by Luminata Forte Western HRP Substrate (Millipore) and images were acquired by Amersham Imager 600 (GE Healthcare).

### Immunoprecipitation

BmN4 cells were resuspended in buffer A [25 mM Tris-HCl (pH 7.4), 150 mM NaCl, 1.5 mM MgCl_2_, 0.15% Triton X-100, 0.5 mM DTT, 1 × Complete EDTA-free protease inhibitor (Roche), 1 × PhosSTOP (Roche)] and incubated on ice for 20 min. The cell suspension was centrifuged at 17,000 × g for 30 min at 4°C. The supernatant was incubated with Dynabeads Protein G (Thermo Fisher/Invitrogen) pre-conjugated with anti-FLAG antibody (M2, Sigma). For HA-immunoprecipitation, the supernatant was pre-incubated with anti-HA antibody (3F10, Sigma) at 4°C for 30 min, and then Dynabeads Protein G (Thermo Fisher/Invitrogen) was added. After incubation at 4°C for 1 h, the beads were washed with buffer B [25 mM Tris-HCl (pH 7.4), 150 mM NaCl, 1.5 mM MgCl_2_, 0.5% Triton X-100, 0.5 mM DTT] five times, and the immunopurified complexes were eluted with 3X FLAG peptide (Sigma) or HA peptide (MBL).

### Purification of recombinant proteins

pCold-Gtsf1-7His or Gtsf1L-6His was transformed into Rosetta 2(DE3) competent cells (Novagen). The cells were cultured at 37°C until the OD_600_ reaches ∼0.6, and then cooled on ice for 30 min. Protein expression was induced with 1 mM IPTG at 15°C for 15 h. The cell pellets were resuspended in TALON lysis buffer [50 mM Hepes-KOH (pH 7.4), 300 mM NaCl, 15 mM imidazole, 0.2 mM TECP, 10 μg/ml leupeptin, 10 μg/ml aprotinin, 1 μg/ml pepstatin A, 1 × EDTA-free protease inhibitor cocktail (Roche)] and sonicated with Bioruptor II (CosmoBio). The cell lysate was centrifuged at 17,000 × g at 4°C for 20 min. The cleared lysate was added to TALON CellThru Resin (Takara) and incubated at 4°C for 1h. After washing the resin with TALON lysis buffer, the bound proteins were eluted with elution buffer [50 mM Hepes-KOH (pH 7.4), 150 mM NaCl, 300 mM imidazole, 0.2 mM TECP]. Peak fractions were pooled and dialyzed with PBS overnight.

### *In vitro* cleavage assay

For Siwi and BmAgo3 immunoprecipitation, BmN4 cells were resuspended in buffer C [25 mM Tris-HCl (pH 7.4), 150 mM NaCl, 1.5 mM MgCl_2_, 1% Triton X-100, 0.5 mM DTT, 1 × Complete EDTA-free protease inhibitor (Roche), 1 × PhosSTOP (Roche)] and incubated on ice for 20 min. The cell suspension was centrifuged at 17,000 × g for 30 min at 4°C. The supernatant was pre-incubated with anti-Siwi or anti-BmAgo3 antibody at 4°C for 1h, and then Dynabeads Protein G (Thermo Fisher/Invitrogen) was added. The mixture was further incubated at 4°C overnight. The beads were washed with buffer C five times, rinsed with buffer D [30 mM Hepes-KOH (pH 7.4), 100 mM KOAc, 2 mM Mg(OAc)_2_, 0.5 mM DTT], and divided for different target cleavage conditions. Target cleavage assay was performed at 28°C for 2 h in a 10 μl reaction containing 3 μl of 40 × reaction mix (Haley *et al*, 2003), 100 nM recombinant proteins and 1 nM ^32^P cap-radiolabeled target RNA. The target RNA was purified by EtOH precipitation following proteinase K treatment and run on 8% denaturing polyacrylamide gel. The images were acquired with the FLA-7000 imaging system (Fujifilm Life Sciences) and quantified using ImageJ/Fiji. The target RNA against an abundant endogenous Siwi or BmAgo3 piRNA was prepared by *in vitro* transcription as described previously (Yoda *et al*, 2010). Primers used for target RNA preparation were described in Table EV1.

### RNAi in silkworm embryos

siRNA-mediated embryonic knockdown was performed as previously described (Kiuchi & Katsuma, 2022; Kiuchi *et al*, 2014). In brief, ∼5□nl of 100□µM siRNA (FASMAC) was injected into a *Bombyx mori* N4 strain embryo within 3□h after oviposition using IM 300 Microinjector (Narishige). The embryos were incubated at 25°C in a humidified petri dish for 120 h. Approximately 12 embryos were collected as one sample and total RNA was extracted. siRNA sequences are listed in Table EV1.

### RNA extraction, and quantitative real-time PCR

Total RNAs prepared by TRI Reagent (Molecular Research Center) were used for real-time PCR and preparation of small RNA libraries. Five hundred nanogram of total RNA was reverse transcribed by PrimeScript RT reagent kit with gDNA eraser (Takara), and qRT-PCR was performed using KAPA SYBR FAST Master Mix (2X) Universal (Kapa Biosystems) and the Thermal Cycler Dice Real Time System (Takara). The primer sequences for real-time PCR are listed in Table EV1.

### Immunofluorescence

BmN4 cells were fixed with 4% paraformaldehyde in PBS at room temperature for 10 min, then the cells were permeabilized with 0.3% Triton X-100 in PBS for 5 min. After pre-incubation with blocking buffer [PBS supplemented with 1% BSA (Sigma) and 0.1% Triton X-100] at room temperature for 1 h, the cells were incubated with primary antibodies [anti-FLAG antibody (M2, Sigma, 1/250), anti-HA antibody (3F10, Roche, 1/300), anti-Siwi antibody (1/400), anti-BmAgo3 antibody (1/250)] in blocking buffer at 4 °C overnight. Alexa Fluor 488 donkey anti-mouse IgG, Alexa Fluor 488 donkey anti-rat IgG, Alexa Fluor 647 donkey anti-rabbit IgG, and Alexa Fluor 647 goat anti-rat IgG secondary antibodies (Thermo Fisher/Invitrogen) were used for detection. For Fig EV2, BmAgo3 was probed with biotinylated anti-BmAgo3 antibody after incubation with the secondary antibody, and the BmAgo3 signal was detected by Cy3-Streptavidin (Jackson ImmunoResearch). Images were captured using Olympus FV3000 confocal laser scanning system with a × 60 oil immersion objective lens (PLAPON 60XO, NA 1.42, Olympus) and processed FV31S-SW Viewer software and Adobe Photoshop Elements 10.

### Small RNA library preparation

Small RNA libraries were constructed from 20−50 nt total RNAs according to the Zamore lab’s open protocol (https://www.dropbox.com/s/r5d7aj3hhyaborq/) with some modifications (Fu *et al*, 2018). Ten synthetic 22-nt oligoribonucleotides (GeneDesign) (Williams *et al*, 2013) were spiked in all samples for normalization. The 3′ adapter was conjugated with an amino CA linker instead of dCC at the 3′ end (GeneDesign) and adenylated using 5′ DNA adenylation kit at the 5′ end (NEB). To reduce a ligation bias, four random nucleotides were included in the 3′ and 5′ adapters [(5′-rAppNNNNTGGAATTCTCGGGTGCCAAGG/amino CA linker-3′) and (5′-GUUCAGAGUUCUACAGUCCGACGAUCNNNN-3′)] and the adapter ligation was performed in the presence of 20% PEG-8000. After the 3′ adapter ligation at 16 °C for ≥16 h, RNAs were size-selected by urea PAGE. For RNA extraction from a polyacrylamide gel, ZR small-RNA PAGE Recovery Kit (ZYMO Research) was used. Small RNA libraries were sequenced using the Illumina HiSeq 4000 platform to obtain 50-nt single-end reads.

### Sequence analysis of small RNA library

After removal of the adaptor sequences by cutadapt (Martin, 2011), 20−45 nt reads without any ambiguous bases were used for the following analyses. These piRNA libraries were normalized by the read count of spikes.

#### Definition of 1U and 10A piRNAs

The RPM of each piRNA was calculated based on an assumption that if 1–26 nt sequences were identical, they were the same piRNA. For piRNA analysis of BmN4 cells and embryos, only highly expressed piRNAs (RPM > 1) in any of the three BmN4 libraries and six embryo libraries were extracted and used. The common parts (1–26 nt) of these piRNAs were mapped to transposons allowing two mismatches with Bowtie (Osanai-Futahashi *et al*, 2008; Langmead *et al*, 2009). The mapped piRNAs, including 16,734 species of 1U (but not 10A) piRNAs and 6,869 species of 10A (but not 1U) piRNAs for BmN4 cell libraries, and 22,374 species of 1U (but not 10A) piRNAs and 8,533 species of 10A (but not 1U) piRNAs for embryo libraries, were extracted and used. Sam files were converted to bam files by SAMtools (Li *et al*, 2009b) and then to bed files by BEDTools (Quinlan & Hall, 2010). The length and 5′-end position for each piRNA were obtained from the bed files using custom R programs.

#### Analysis of Fem and Masc piRNAs

Each piRNA library was mapped to sequences of *Fem* RNA and *Masc* mRNA (Kiuchi *et al*, 2014) with Bowtie (Langmead *et al*, 2009) allowing one mismatch. Sam files were converted to bam files by SAMtools (Li *et al*, 2009b) and then to bed files by BEDTools (Quinlan & Hall, 2010). The length and 5′-end position for each piRNA were obtained from the bed files using custom R programs.

#### Calculation of 1U/10A-strand bias per transposon

1U/10A-strand bias was calculated for 750 transposons with RPM greater than 10 for both the sense and antisense strands in the siGFP#1 library of embryos. For each transposon, the 1U bias and 10A bias for each strand were calculated, and then the strand with superior 1U bias (1U strand) and the strand with superior 10A bias (10A strand) were defined. As a result, 330 transposons had more 1U (but not 10A) piRNAs in the sense strand and 345 transposons had more 1U (but not 10A) piRNAs in the antisense strand. The 1U strand piRNA/10A strand piRNA of each transposon was then calculated for each small RNA library, and the z-scores were calculated by the scale function of R. The “fields” package of R was used to plot the heatmaps.

#### Calculation of ping-pong Z-score

The calculation of Z-score was based on a previous study (Zhang *et al*, 2011), with some modifications. The ping-pong Z-score is the difference between the mean of the scores for each overlap from nucleotides 1 to 21 divided by the standard deviation of the background distance scores, defined as the distances 1–9 and 11–21. The calculation of the Z-scores was performed separately for 1U (but not 10A) piRNAs and 10A (but not 1U) piRNAs on the sense strands of transposons.

### Prediction of the Gtsf-PIWI complexes

Protein structures were predicted using AlphaFold ver. 2.1.2 (Jumper *et al*, 2021; Evans *et al*, 2021) installed on a local computer via Docker and CUDA Toolkit 11.1. full_dbs was used as a database and max_template_date was defined as 2021-07-14. Two protein sequences and a multimer model were used to predict protein complexes (Evans *et al*, 2021). The heatmaps of predicted aligned errors were made using the pae2png.py script (https://github.com/CYP152N1/plddt2csv). Visualization of the predicted structure was done using PyMOL.

## Expanded View Figure Legends

**Figure EV1.**
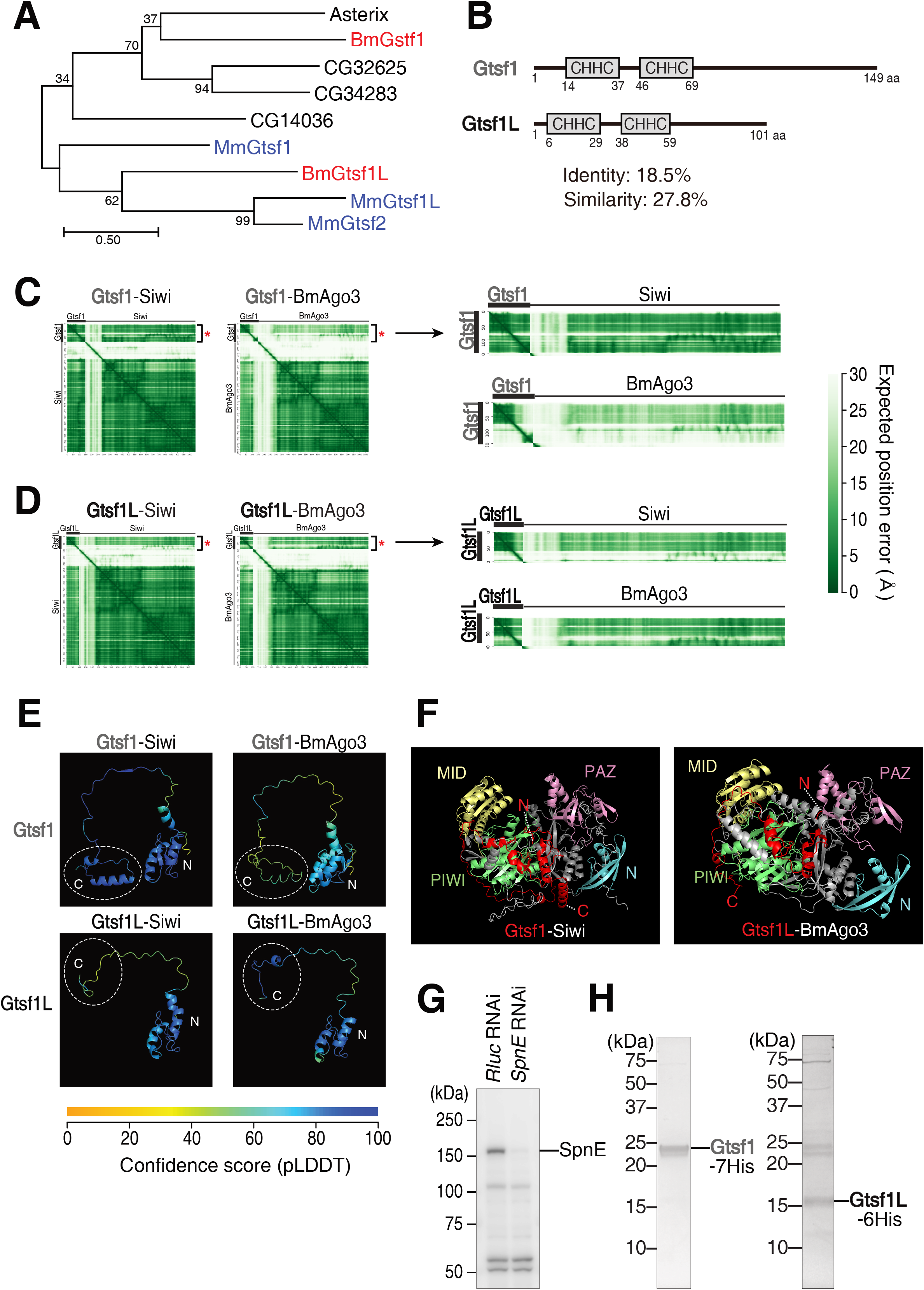
Prediction of Gtsf-PIWI complexes using AlphaFold-Multimer. **A** Phylogenetic tree of fly (black), mouse (blue), and silkworm (red) Gtsf homologs. The phylogenetic tree was constructed using the maximum likelihood method with bootstrapping in MEGAX (Tamura *et al*, 2021). The numbers at the branch point indicate the bootstrap support values. **B** Schematic presentation of the domain structure of *Bombyx* Gtsf1 and Gtsf1L. **C, D** Predicted Aligned Error (PAE) plots for the top-ranked model of Gtsf1-PIWI (C) and Gtsf1L-PIWI (D) complexes predicted by AlphaFold-Multimer. The regions indicated with asterisks are enlarged on the right. **E** AlphaFold-Multimer-predicted structure of Gtsf1 and Gtsf1L in complex with Siwi or BmAgo3. The C-terminal structure (white dotted circles) of Gtsf1 and Gtsf1L is predicted with higher confidence in complex with Siwi and BmAgo3, respectively. **F** AlphaFold-Multimer-predicted structure of the Gtsf1-Siwi and Gtsf1L-BmAgo3 complexes. The domain definition (the N, PAZ, MID, and PIWI domains) of Siwi follows a previous study (Matsumoto *et al*, 2016), and the corresponding regions of BmAgo3 based on the sequence alignment with Siwi are colored. **G** Western blot analysis of whole cell lysate of BmN4 cells treated with dsRNA for *Rluc* (control) or *SpnE*. The anti-SpnE antibody successfully detects endogenous SpnE. **H** CBB staining of purified Gtsf1 and Gtsf1L proteins used for *in vitro* target cleavage assay.

**Figure EV2.**
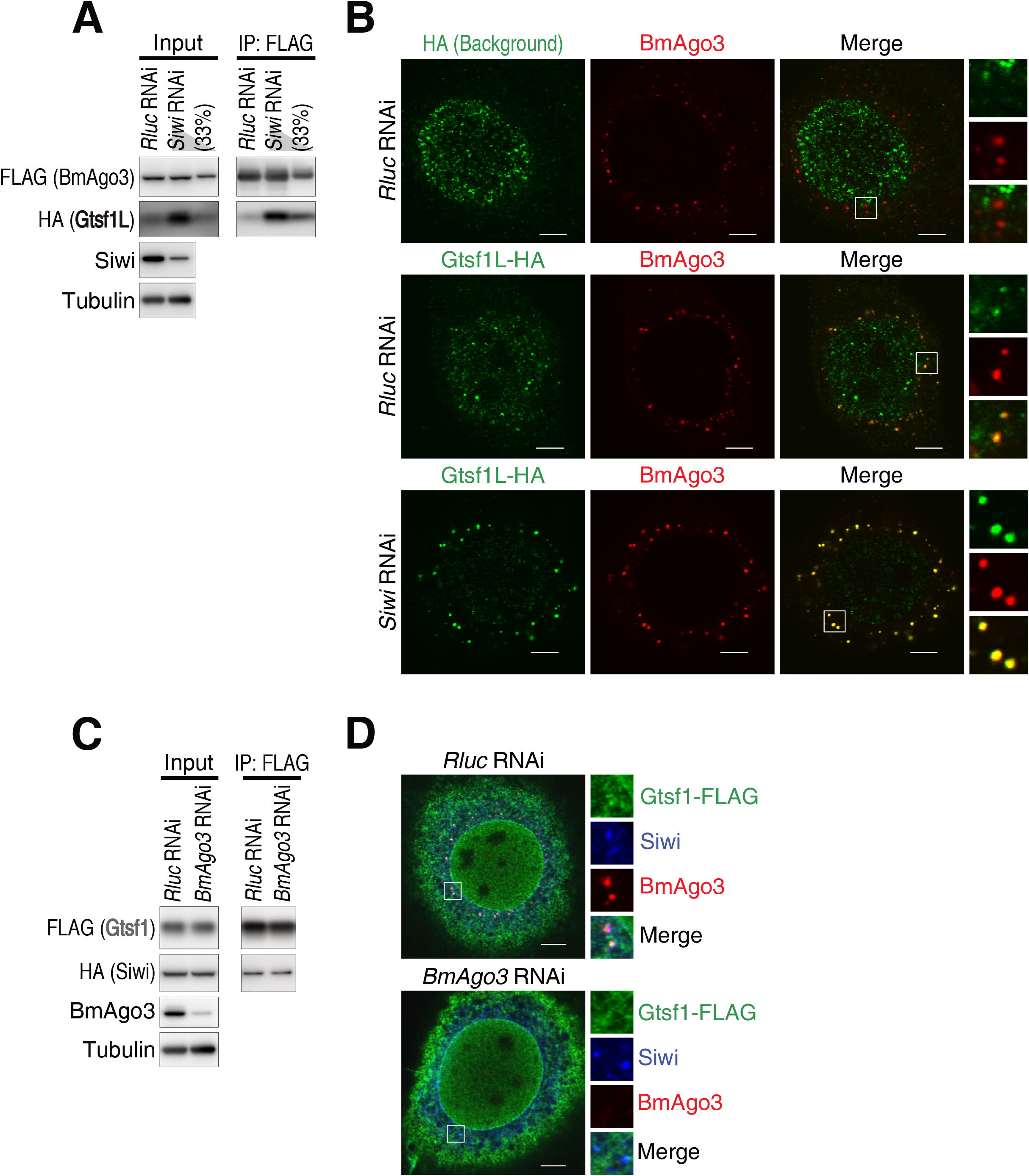
*Siwi* knockdown stabilizes the expression of Gtsf1L and increases the association of Gtsf1L with BmAgo3. **A** Western blot analysis of the interaction between Gtsf1L-HA and FLAG-BmAgo3 in *Siwi* knockdown. **B** Subcellular localization of Gtsf1L-HA and BmAgo3 in BmN4 cells transfected with dsRNA for *Rluc* (control) or *Siwi*. Gtsf1 localizes to BmAgo3 granules (middle panel) and this is strongly enhanced by *Siwi* knockdown (lower panel). The nuclear signal of Gtsf1L-HA is nonspecific staining of anti-HA antibody (upper panel). Scale bar, 5 μm. **C** Western blot analysis of the interaction between Gtsf1-FLAG and HA-Siwi in *BmAgo3* knockdown. The interaction between Gtsf1 and Siwi is unaffected by *BmAgo3* knockdown. **D** Subcellular localization of Gtsf1-FLAG and Siwi in BmN4 cells transfected with dsRNA for *Rluc* (control) or *BmAgo3*. No accumulation of Gtsf1 in Siwi granules is detected in *BmAgo3* knockdown. Scale bar, 5 μm.

**Figure EV3.**
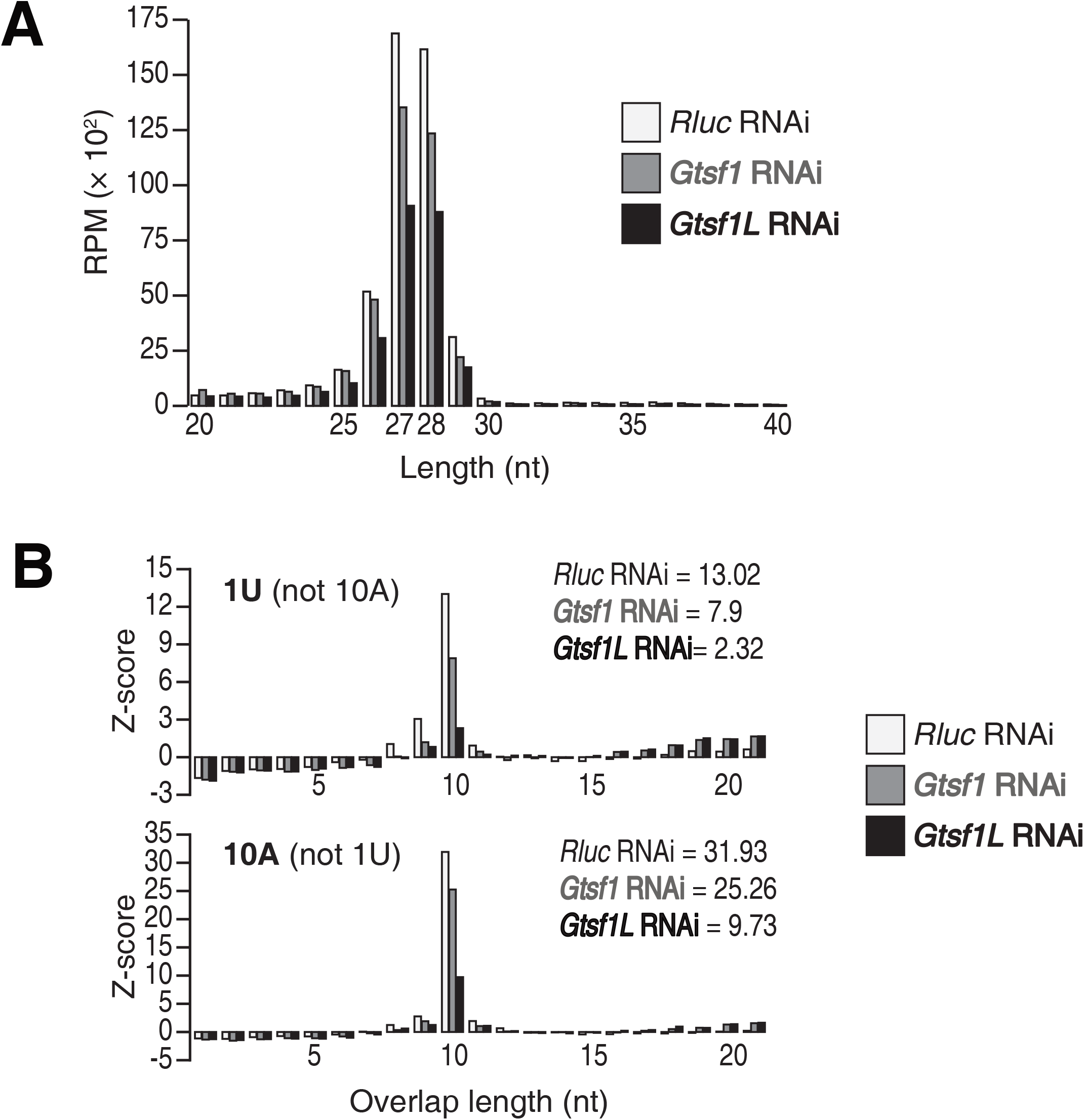
Knockdown of *Gtsf1* or *Gtsf1L* decreases piRNA levels without affecting their length. **A** Length distribution of total piRNAs in *Gtsf1* or *Gtsf1L* knockdown. Both *Gtsf1* and *Gtsf1L* knockdown modestly decrease mature piRNA levels. **B** The Z-scores of 5’-5’ overlapped piRNAs (y-axis) at the indicated length (x-axis) for 1U not 10A (upper) and 10A not 1U (lower) piRNAs in the small RNA libraries from BmN4 cells transfected with the indicated dsRNAs. The Z-scores at the 10 nt overlap (ping-pong signature) are shown in the top-right of the graphs.

**Figure EV4.**
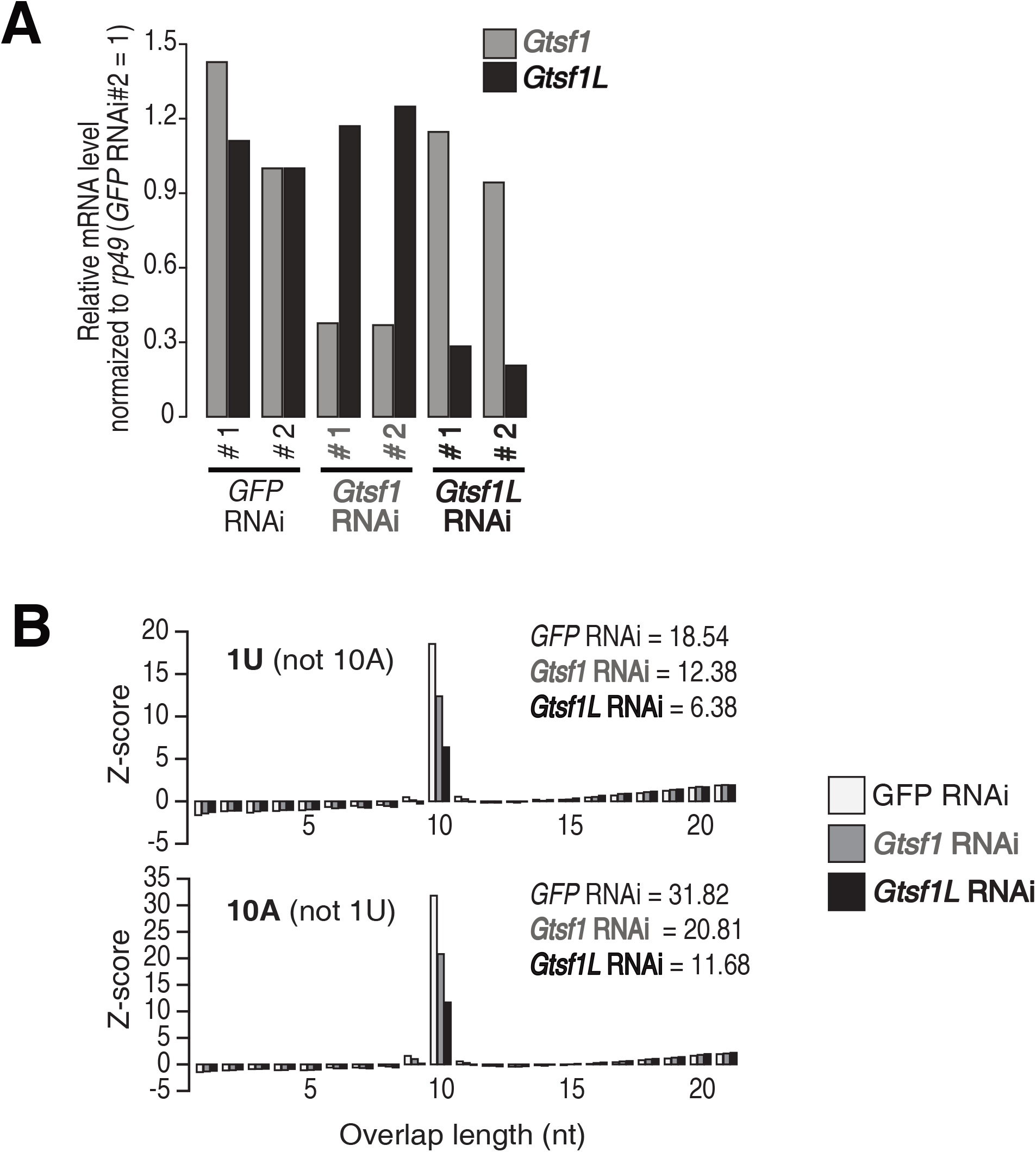
*Gtsf1*- and *Gtsf1L*-knockdown in silkworm embryos. **A** Quantitative real-time PCR analysis of the expression of *Gtsf1* and *Gtsf1L* in siRNA-injected embryos. The embryos were analyzed 120 hours post-injection. *GFP* siRNA was used as a control and the values were normalized to that of *rp49*. *Gtsf1* and *Gtsf1L* mRNAs were specifically decreased by each siRNA. **B** The Z-scores of 5’-5’ overlapped piRNAs (y-axis) at the indicated length (x-axis) for 1U not 10A (upper) and 10A not 1U (lower) piRNAs in the small RNA libraries from embryos injected with the indicated siRNAs. The Z-scores at the 10 nt overlap (ping-pong signature) are shown in the top-right of the graphs.

## Notes

### Competing Interest Statement

The authors have declared no competing interest.

### Summary of Updates

Minor changes in the text and Figure 3, EV3, and EV4.

